# Biotinylation by antibody recognition - A novel method for proximity labeling

**DOI:** 10.1101/069187

**Authors:** Daniel Z Bar, Kathleen Atkatsh, Urraca Tavarez, Michael R Erdos, Yosef Gruenbaum, Francis S Collins

## Abstract

Identification of protein-protein interactions is a major goal of biological research. Despite technical advances over the last two decades, important but still largely unsolved challenges include the high-throughput detection of interactions directly from primary tissue and the identification of interactors of insoluble proteins that form higher-order structures. We have developed a novel, proximity-based labeling approach that uses antibodies to guide biotin deposition onto adjacent proteins in fixed cells and primary tissues. We used this method to profile the dynamic interactome of lamin A/C in multiple cell and tissue types under various treatment conditions. Our results suggest a considerable variation in the composition of the nuclear envelope of different tissues. Of note, DNA damage response proteins Ku70 and Ku80 are more abundant in the vicinity of lamin A/C after thermal stress. This increased affinity also applies to the progerin isoform, potentially contributing to the premature aging phenotype of Hutchinson-Gilford progeria syndrome. The ability to detect protein-protein interactions in intact tissues, and to compare affinities quantitatively under different conditions or in the presence of disease mutations, can provide a new window into cell biology and disease pathogenesis.

## Introduction

Protein-protein interactions (PPI) are critical to the function of all living cells. The protein interactome is dynamic: interactions may change with time, developmental stage, cell cycle progression, or tissue type. Proteins often act together; but depending on their binding partners, individual proteins can produce variable and even opposite biological effects. Characterizing PPI can thus provide important information about the locations and functions of a protein of interest. Alterations of specific PPI, for instance by non-synonymous mutations, may correspondingly alter an organism′s phenotype. However, the effect of specific mutations on tissue-specific protein interactomes has rarely been studied in any real detail.

Distinct mutations within a single gene may result in a plethora of diseases. Over 400 different mutations of the lamin A/C (LMNA) gene give rise to more than 14 distinct phenotypes that affect a variety of tissues -- among them lipodystrophies, muscular dystrophies, peripheral neuropathies, and premature aging syndromes. The mechanistic links between these single-gene mutations and their highly variable phenotypes remain to be elucidated. Tissue-specific interactors may offer an explanation, but characterization of the tissue-specific lamin interactome is challenging and has yet to be accomplished.

Lamins are type V nuclear intermediate filaments that localize primarily to the nuclear envelope (NE). Analysis of the lamin interactome by standard Co-Immunoprecipitation (CoIP) is impractical because the insoluble high order structures created by the lamin filaments result in superfluous precipitation of unrelated proteins. Several attempts to identify lamin interactors and to characterize the protein composition of the NE have been made with various genetic and biochemical approaches (abbreviated BioID^1^, OneSTrEP-tag^2^, Y2H^3^, Lamin A/C tail^4^, Liver LB BioID^5^ and FACS + salt^6^). While multiple known and novel interactors were identified by these methods, their applications were limited to single cell lines and showed only a modest overlap amongst different datasets (see below). Additionally, these methods cannot be directly applied to existing animal models and to primary human tissue.

Here we describe the development and validation of a novel proximity-based labeling method for detecting potential protein-protein interactions.

Previously published proximity-based labeling methods rely on a deposition of a tag, typically biotin, on molecules adjacent to a target of interest. For example, fusion of a promiscuous biotin ligase to the lamin A/C gene was used to identify proteins proximal to the lamin A/C protein in transfected cells^1^. To improve the chemistry of the reaction, Ting et al.^7^ fused a peroxidase to proteins of interest and used phenol-biotin to successfully identify mitochondrial proteins. While these strategies substantially outperform traditional co-immunoprecipitation methods, they have considerable limitations. Most significantly, these methods require the prior insertion of a fusion gene in cell lines or in the animal germline, and thus cannot be used on primary human tissue samples or in most existing animal models. These methods also do not allow for the selective detection of proteins carrying a specific post-translational modification.

To overcome these limitations, we have developed an antibody-guided, proximity-based labeling method denoted Biotinylation by Antibody Recognition (BAR) and applied this method to identify the interactors of lamin A/C in immortalized cell lines, primary cell culture, and primary human muscle and adipose tissues. We further expanded the method to include differential proteomics and identified stress and mutation-induced changes to the composition of the NE. Finally, we detected interactions between progerin and DNA damage response proteins that may play a role in the phenotype of Hutchinson-Gilford progeria syndrome.

## Results

We developed and applied a novel method, termed Biotinylation by Antibody Recognition (BAR), to identify proteins in the vicinity of an antigen (Fig. 1A). In a fixed and permeabilized tissue sample, a primary antibody is used to target a protein of interest. In the presence of hyd\rogen peroxide and phenol biotin, a secondary HRP-conjugated antibody creates free radicals, resulting in a biotinylation of proteins in close proximity to the target protein. As biotin is covalently attached, harsh conditions may then be used for reverse cross-linking and protein solubilization. Streptavidin-coated beads are used to immunoprecipitate the biotinylated proteins, which are then detected by tandem mass-spectrometry.

**Figure 1:**
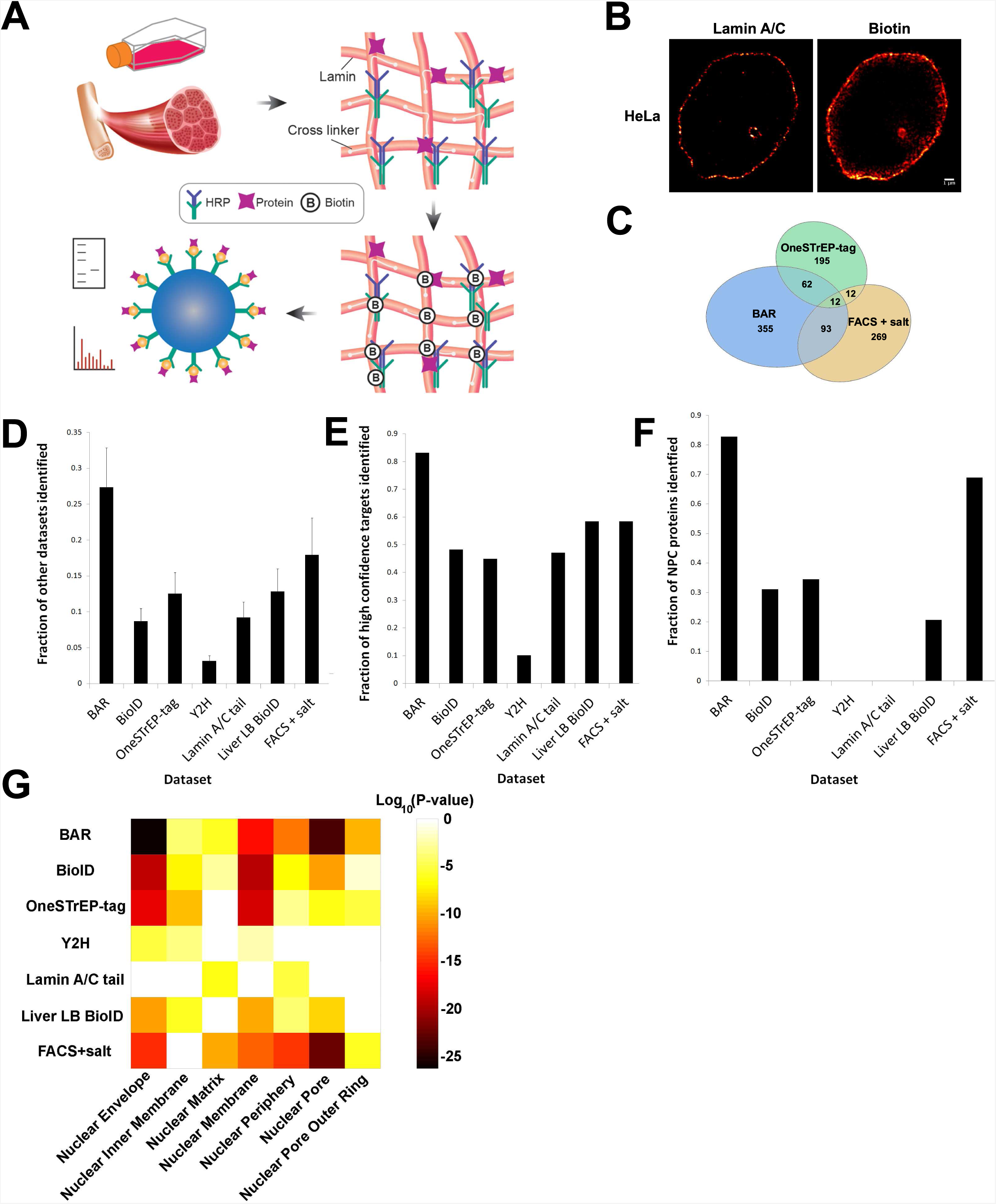
Biotinylation by antibody recognition. (**A**) Schematic representation of the method, applied in this case to laminA/C. (**B**) Super-resolution microscopy showing biotin deposition in the vicinity of the NE, confirming the tight spatial resolution of the protein labeling method. (**C**) Venn diagram showing the larger overlap of the ratiometric on-bead labeling dataset (BAR) with the two largest datasets compared. (**D**) For each of the 7 datasets (our ratio-metric on-bead dataset, as well as 6 other published datasets), the average fraction of all other datasets covered by it is shown. (**E**) The fraction of high confidence targets identified in each dataset is shown. (**F**) The fraction of 29 known nuclear core complex roteins identified in each dataset is shown. (**G**) Heat map showing P-values for 7 Gene Ontology cellular component categories across our ratiometric on-bead dataset, as well as 6 other published datasets. P-values were calculated using PANTHER overrepresentation test^8^

### Lamin A/C proximity labeling in HeLa cells

We applied BAR to identify proteins in the vicinity of lamin A/C in HeLa cells. The labeling radius was limited by reducing the reaction time and selecting compatible blocking reagents (see Supplementary Methods). Other attempts to reduce the labeling radius, such as altering temperature or viscosity, decreased the signal to noise ratio (see Supplementary Methods). Biotin was successfully deposited at the NE, as evident from super-resolution microscopy (Fig. 1B). Biotinylated proteins were pulled down with streptavidin-coated beads and analysed by Western blot and mass-spectrometry of in-gel digested proteins (Supplementary Dataset, Supplementary Fig. 1A-C). In addition to identifying the targeted lamin A/C, we identified multiple known interactors of lamin A/C, including lamin B1, lamin B2 and lamina-associated polypeptide 2 (LAP2) (Supplementary Fig. 1B, Supplementary Dataset). These proteins were either found exclusively or enriched in the sample as compared to the control (Supplementary Fig. 1C, Supplementary Dataset).

### Ratiometric analysis

To improve on these results, we sought to increase signal intensity. Reaction time determines the labeling radius and signal intensity. While limiting the labeling radius reduced the number of non-NE proteins identified, it also decreased the signal intensity both for rare and abundant NE proteins (Supplementary Fig. 1D, Supplementary Dataset). By contrast, increasing the reaction time results in signal arising from leakage to non-nuclear envelope proteins. To overcome this, we employed a ratiometric based *enrichment score*. An adjacent control can occasionally be found for some subcellular locations ^9^, however this may not be the case for proteins with multiple subcellular locations or proteins found between distinct cellular structures. Thus, we determined enrichment by comparing the streptavidin-bound fraction with the unbound fraction. For proteins in proximity to our target, biotin labeling will drive enrichment in the bound over unbound fraction. By contrast, as the signal from the NE decays with distance, abundant proteins that are labeled due to signal leakage will be underrepresented in the bound compared to the unbound fraction. The enrichment score was calculated as the normalized bound to unbound protein area ratio or the bound to no antibody control, the lesser of the two. Finally, we replaced in-gel digestion with an on-bead digestion, reducing handling time and variance, as well as mass-spectrometry time and cost. Overall, compared to our “in-gel” dataset, ratiometric labeling resulted in a modest improvement in dataset quality (see below) and a significant improvement in protein signal and peptide count, thus enabling better quantification (Supplementary Dataset).

### Comparison with previous attempts to characterize the lamin interactome

We compared our BAR results to that obtained from six other methods. These datasets were generated by various biochemical and genetic methods applied to different cell lines and yeast. Each dataset is composed of true and false positive results. We calculated the average fraction of all other datasets covered by a specific dataset. Because true positives are more likely to be enriched by multiple independent experiments, a higher percentage overlap is indicative of higher dataset quality. The overlap between the six collected datasets, while statistically significant, is small (Fig. 1C,D) even if we account for the different cell lines used (see HeLa and HepG2 comparison in Supplementary Dataset). Datasets from all other methods had an average overlap of 11% (3-17% range) with other lamin interactome datasets (Supplementary Fig. 1E-G, Fig. 1D). By contrast, 24% and 27% of previously identified putative lamin binding proteins were identified by our BAR “in-gel” digestion and ratiometric experiments, respectively (Supplementary Fig. 1F and Fig. 1D). To gain a better estimate of coverage of true positive results, we defined high confidence interactors as proteins identified by three or more datasets. Of these 81 proteins, 81% (66/81) were identified by our “in-gel” dataset, compared with 45% (37/81) proteins on average and a maximum of 64% (52/81) by other methods (Supplementary Fig. 1G). Within our ratiometric labeling dataset, 83% (74/89) high confidence proteins were detected, compared with an average of 45% (40/89) and a maximum of 58% (52/89) for all other datasets (Fig. 1E). Similarly, 83% (24/29) of the nuclear pore complex proteins, found in close proximity to lamin A/C, were identified by ratiometric labeling (Fig. 1F). Similar conclusions resulted from comparing the Gene Ontology enrichments of the various datasets (Fig. 1G).

### Lamin A/C proximity labeling in primary human tissue

An underlying assumption of cell culture usage is that it mimics to a significant extent relevant processes that occur throughout the entire organism. While this assumption is often valid, in many cases it is not. To overcome such limitations, we attempted to identify lamin A/C interactors in primary human tissue. As laminopathies often affect fat distribution and cause muscle weakness, we analyzed human skeletal muscle, smooth muscle, and adipose samples obtained post-mortem. We successfully labeled the NE in post-mortem tissue samples. Nuclear envelope morphology in these samples deviated considerably from the classical view of a smooth round sphere around the nucleus. For example, muscle myofibrils created grooves in the nucleus, which dictated not only the distribution of lamin A/C, but also of DNA (Fig. 2A, Supplementary Fig. 2A). Some nuclei in adipose had a doughnut shape (Fig. 2B), a phenotype that was previously reported in cell culture but not in primary human tissues ^10^. While different tissues had multiple lamin A/C interactors in common, some proteins were either found exclusively or with greater abundance in a particular tissue. For example, nesprin proteins (SYNE1 and SYNE2) are known lamin A/C interactors and members of the Linker of Nucleoskeleton and Cytoskeleton (LINC) complex^11,12^. In HeLa cells, these proteins were either detected with a low peptide count of one to two peptides or not detected at all. By contrast, nesprins were identified in all our human muscle samples with a unique peptide count of up to 14 (Supplementary Dataset). Similarly, Muscular LMNA-interacting protein (MLIP) was found in all muscle samples but not in HeLa or adipose samples (Supplementary Dataset). Proteins identified by two or more unique peptides in one experiment had a ∼90% chance of being detected (at any level) in a replicate experiment (Fig. 2C). However, signal correlation between replications, while highly statistically significant, made quantification of small differences without multiple repetition impossible (Fig. 2D, see Differential proteomics below). Overall, skeletal muscle tissue showed higher similarity to smooth muscle tissue than to adipose tissue. Of 141 skeletal muscle NE proteins identified with two or more peptides, 93.6% (132/141) were also identified in smooth muscles, but only 42.6% (60/141) were found in adipose tissue (Fig. 2E). These results suggest that the NE composition varies significantly among different tissues.

**Figure 2:**
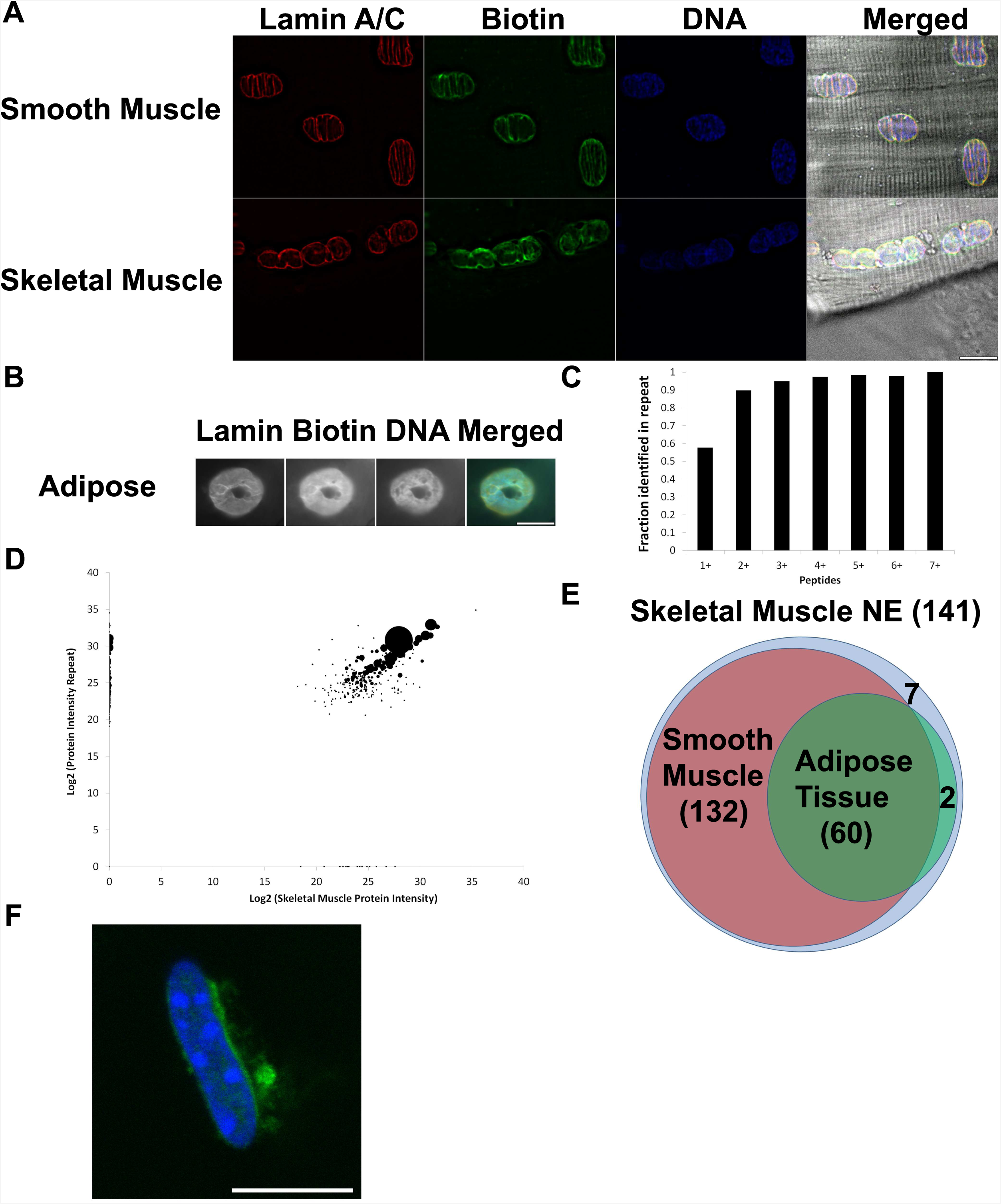
Identification of NE proteins in primary human tissue. (**A**) Imaging of primary human skeletal and smooth muscle tissue. (**B**) Doughnut shaped nuclei from primary human adipose tissue. (**C**) Fraction of proteins with enrichment score >1 identified in a repeated human skeletal muscle BAR experiment, as a function of unique peptides identified. (**D**) Signal correlation between proteins having an enrichment score > 1 in the two skeletal muscle samples. Dot size indicates minimal number of unique peptides identified in either of the experiments. Proteins detected in only one sample were set to zero at the other. Large dot is Titin. (**E**) A venn diagram showing the number of skeletal muscle proteins identified by at least 3 peptides having an enrichment score >1, and how many of these proteins were identified (without any filtering criteria) in smooth muscle and adipose tissue. Diagram not to scale. (**F**) SGCA localizes to the nuclear periphery in mouse skeletal muscle. Scale bar - 10 µm.

Muscle tissues showed NE enrichment for several proteins mutated in muscular dystrophies and cardiomyopathies, including multiple members of the dystrophin-glycoprotein complex (Supplementary Dataset). While members of this complex were not previously reported to be associated with lamin A/C, fluorescent protein fusion and antibody staining in cell culture^13^ suggests that this complex is enriched at close proximity to the NE, presumably in the perinuclear ER. Indeed, we were able to replicate these results in primary human and mouse skeletal muscle tissue (Fig. 2F and Supplementary Fig. 2B). To gain further support for this association, we applied BAR to identify proteins in close proximity to α-sarcoglycan (SGCA), a member of the dystrophin-glycoprotein complex. We successfully identified multiple members of the dystrophin-glycoprotein complex (Supplementary Dataset). In support of our hypothesis, lamin A/C was also moderately enriched in primary skeletal muscle when BAR was applied with an antibody against SGCA (Supplementary Dataset).

### Differential proteomics

Qualitative and quantitative changes may be observed when comparing the protein interactome under different conditions, such as the presence or absence of a genetic variation, application of a treatment protocol, or variation of a tissue type. Identifying such changes is important for understanding the function of the target protein in that context. While a protein of interest may be in close proximity to multiple other proteins, only a small fraction of those proteins may be relevant to a specific function or process, and this fraction is more likely to change in that context. To expand the usability of our method, we used Stable Isotope Labeling by Amino Acids in Cell Culture (SILAC) to perform differential proteomics. Without treatment, NE proteins extracted from cells grown side-by-side on either heavy or light medium and analyzed by mass-spectrometry showed minimal variation (Fig. 3A). As a proof of concept for differential proteomics, we compared the NE composition of naive HeLa cells with cells subjected to a 2-hour 43°C heat shock. As expected, in control (naive) cells, the heavy to light ratio was very close to one, particularly when multiple peptides were used to calculate the ratio (Fig. 3B). By contrast, when comparing a control with the shocked sample, multiple proteins displayed deviant heavy to light ratios (Fig. 3C - blue bars). These changes were mirrored in a reciprocal experiment (Fig. 3C - red bars). Proteins identified include expected targets, such as multiple members of the heat shock protein family. As these proteins become abundant during heat shock conditions, the NE bound fraction is also expected to increase. By contrast, we also identified elevated levels of non heat-shock proteins. For example, Ku70 and Ku80 (also known as XRCC6 and XRCC5, respectively) are regulators of DNA-PK catalytic subunit (DNA-PKcs, also known as PRKDC) and members of the non-homologous end joining pathway. We found Ku70 and Ku80 to be ∼2.5-fold more abundant near the NE following heat shock (p = 10^−5^ and 0.0005 for Ku70. p = 0.007 and 0.002 for Ku80). In contrast, no enrichment was seen for DNA-PKcs. We validated the conclusion that Ku70 and Ku80 localize near lamin A/C using Förster resonance energy transfer (Supplementary Fig. 3) and also observed an increase in Ku70 nuclear peripheral localization using immunofluorescence following heat shock (Fig. 3D). Ku70 was previously identified^1,2^ as a lamin A/C binding protein and heat shock inactivates Ku80 and drives its aggregation^14^. HSPA8, known to bind Ku70 and suppress its nuclear import, also showed a similar NE increase (p < 10^−6^ for each of the two experiments). To confirm that these changes reflect changes in molecular localization, and not total protein abundance, we visualized Ku70 and Ku80 by imaging HeLa cells expressing GFP-Ku70 and GFP-Ku80 during heat shock. As expected, total protein level did not increase during a 2 hour heat shock (Supplementary Fig. 4).

**Figure 3:**
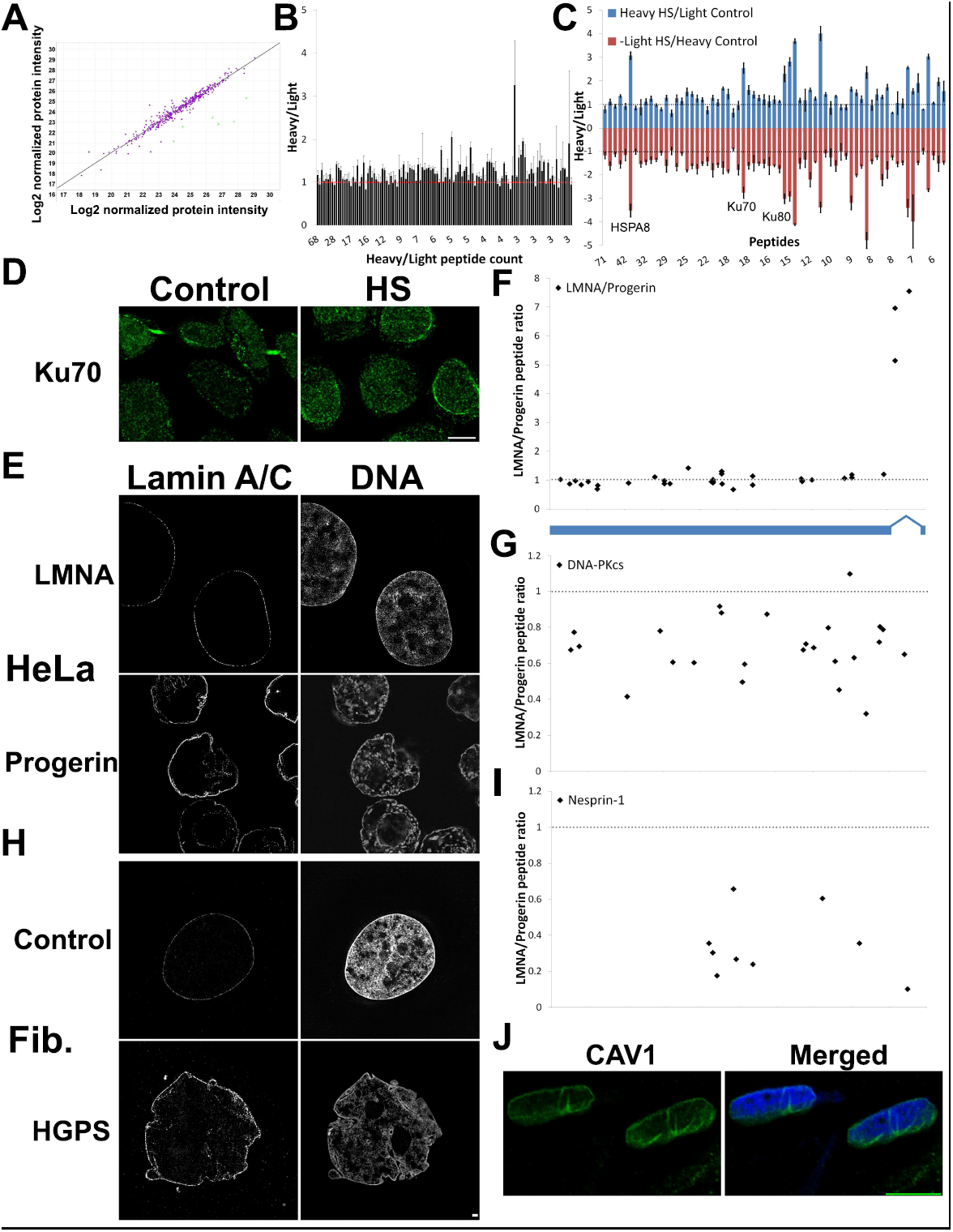
Differential proteomics. (**A**) Scatter plot of BAR extracted proteins from heavy vs. light labeled untreated HeLa cells. (**B**) Heavy/Light ratio of HeLa NE proteins in untreated cells. Proteins are ordered by peptide count. Proteins passing the filtering criteria in the ratiometric on-bead dataset were considered NE proteins. (**C**) Heavy/light ratio of HeLa NE proteins in untreated versus heat shocked cells. Proteins are ordered by sum of peptides used to calculate the heavy/light ratio. (**D**) Ku70 subcellular localization in HeLa cells, visualized with immunofluorescence, before and after heat shock. Scale bar - 10 μm. (**E**) Structured illumination super-resolution microscopy of HeLa cells transfected with GFP-LMNA or GFP-Progerin. (**F**) Heavy (lamin A/C) to light (progerin) peptide ratio of GFP-LMNA/Progerin in transfected HeLa cells. Progerin protein model (blue) indicates the location of the 50aa deletion, overlapping with the last 3 peptides. (**G**) Heavy (lamin A/C) to light (progerin) DNA-PKcs peptide ratio in GFP-LMNA/Progerin transfected HeLa cells. (**H**) Structured illumination super-resolution microscopy of control and HGPS-derived fibroblasts. Scale bar - 1 μm. (**I**) Nesprin-1 peptide ratio in control (heavy) vs HGPS (light) fibroblasts. (**J**) CAV1 immunofluorescence in primary human muscle tissue. Scale bar - 10 μm.human muscle tissue. Scale bar - 10 μm.

### Progerin proximity labeling in transfected HeLa cells

Roughly 90% of cases of the premature aging disease Hutchinson Gilford progeria syndrome (HGPS) are caused by a *de-novo* synonymous mutation in exon 11 at position 1824 of the lamin A/C gene^15^. This mutation activates a cryptic splice site, resulting in a protein lacking 50 amino acids, termed progerin. During maturation of wild-type lamin A/C, it is farnesylated and carboxymethylated at its C-terminus. However, these modifications are not found in the mature protein, due to the cleavage of the lamin tail by ZMPSTE24. By contrast, progerin lacks the ZMPSTE24 recognition site, resulting in a permanently farnesylated and carboxymethylated protein. To compare the lamin A/C and progerin interactome, we applied BAR to HeLa cells transfected with GFP-LMNA or GFP-progerin and used a GFP antibody to direct biotin labeling. By observing the cells 24 hours after transfection, we were able to detect changes to the composition of the NE resulting directly from the acute expression of progerin, rather than from later byproducts of chronic expression, such as cellular senescence. As expected, the heavy to light ratio for most lamin A/C peptides was close to one. Only three peptides exhibited a significant deviation from this ratio, all of which overlapped with the exon 11-encoded 50 amino acid stretch found only in lamin A/C and not in progerin, thus demonstrating our ability to distinguish between isoforms even in a wild-type lamin A/C endogenous background (Fig. 3F). Among the proteins preferentially binding progerin was DNA-PKcs, previously reported to bind progerin^16^, Fig. 3G). Additionally, we found progerin to preferentially bind Ku70 and Ku80, for which progerin binding was not previously reported (Supplementary Dataset).

### Nuclear envelope proximity labeling in primary HGPS fibroblasts

The NE composition of primary tissue and primary tissue culture differs from that of HeLa cell culture. To detect changes in NE composition resulting from progerin expression, we grew primary control fibroblasts in SILAC medium (Fig. 3H, Supplementary Dataset) and compared the NE composition with fibroblasts donated by a HGPS patient. We detected changes in the abundance of several known NE proteins, including lamin B2, nesprin-1 (Fig. 3I), SUN2 and RanBP2. Additionally, in HGPS fibroblasts we saw a NE decrease in Polymerase I and transcript release factor (PTRF) and its interactor caveolin-1 (CAV1). Mutations in PTRF and Caveolin family member proteins are associated with lipodystrophy and muscular dystrophy^17^. To gain further insight into the roles of CAV1, we immunostained CAV1 in HGPS fibroblasts. While some CAV1 foci were seen at the nuclear periphery (Supplementary Fig. 5), only a small fraction of CAV1 localizes near the nuclear membrane and may be available for NE interactions. Next we looked at a more biologically relevant tissue, primary human skeletal muscle. In this tissue, a clear NE localization with a more smooth distribution was seen, suggesting a significant fraction of CAV1 is found at the nuclear membrane (Fig 3J).

## Discussion

We have developed a novel and generalizable method for the identification of proteins in close proximity to an antigen and successfully applied it to characterize the NE composition in multiple cell lines and tissues. Our results expand previous studies that have characterized the interactions of NE proteins with lamin A/C, and demonstrate how the results of such studies can vary significantly between tissues. We have also compared the interactome of progerin to that of lamin A/C, replicating multiple known differences and identifying novel ones. Finally, we showed that heat shock, a form of proteotoxic stress, but not genotoxic stress, drives Ku70 and Ku80 to the NE (Supplementary Dataset).

### Advantages and Limitations of the Proximity-based Labeling Method

Networks of protein-protein interactions are dominated by weak interactions^18^. While CoIP-based methods are being developed to identify these interactions using ultra-high-throughput proteomics and strong statistical methods^18^, previously reported proximity-based labeling methods outperform CoIP for challenging proteins like lamin A/C, and are likely to be useful in identifications of weak and transient interactions in non-challenging proteins as well. By replacing enzyme fusion with antibodies, our method --while based on a chemistry similar to that used by Ting et al. ^7^--offers several advantages. Unlike previous proximity-based labeling methods, BAR does not need a separate cell line or animal model to be generated for every protein of interest. Our use of antibodies prevents any protein fusion related artifacts. Thus, this is the first method that enables the use of proximity based labeling to directly probe the lamin interactome of primary human tissue. Nonetheless, our method has several drawbacks. These include the requirement for a monospecific antibody that works well in tissue sections, sensitivity to fixation artifacts, and inability of performing *in vivo* labeling.

### Accuracy estimation

To estimate the accuracy of our method, we compared our results with two datasets, a small high-confidence list of nuclear pore complex proteins and a larger list of lamin-binding proteins identified by multiple independent labs using different methodologies. We identified more than 80% of the proteins present in each of the lists (Fig 1E,F). These results compare favorably with previous attempts to characterize the lamin interactome and suggest that robust true interactors are likely to be picked up by our method. However, like most other datasets, we rely on mass-spectrometry to identify the pulled-down proteins. We cannot exclude the possibility that many more low abundant, weak or transient interactors are below the current detection limit of the mass spectrometer. While BAR successfully identified the majority of high confidence proteins, most putative interactors identified by us, as well as by all previous attempts, are not supported by cross-study comparisons. Overall, ∼43% of proteins identified in our HeLa datasets were identified in at least one other dataset, similar to the average for all other datasets compared. This can only partially be accounted for by biological differences between cell lines used. It is likely that all current methods have high false positive and false negative rates. For proximity labeling, false positives may partially result from non-interacting proteins found in relative proximity. In experiments aimed at increasing true positive to false positive ratio at the expense of identification of true positives and signal intensity, some improvement should be achievable by using BAR with new proximity-based labeling chemistries, like *in vivo* proximity labeling (IPL; ref. ^19^), offering the possibility to probe only direct interactions.

### Tissue specificity

Previous studies characterizing the lamin interactome analyzed single cell lines. To gain insight into how specific mutations in lamin A/C cause multiple distinct clinical conditions, we analyzed the lamin interactome of multiple cell lines and primary human tissues. Certain proteins known to be clinically relevant, which were not identified by previous studies nor by BAR as lamin A/C interactors in HeLa cells, were identified by BAR in disease-relevant primary tissues. Primary muscle tissue shows NE enrichment for members of the dystrophin-glycoprotein complex. Mutations in this complex can result in various muscle dystrophies that resemble the muscle dystrophies caused by lamin A/C mutations. We show that multiple members of the dystrophin-glycoprotein complex are found in close proximity to the NE in skeletal muscle (Fig. 2F, Supplementary Fig. 2, Supplementary Dataset). Interestingly, mutations to the dystrophin protein can cause NE abnormalities that progress with cell passage^20^. The dystrophin-glycoprotein complex has a structural role in linking the cytoskeleton to the extracellular matrix. In view of its enrichment at the nuclear periphery (Fig. 2F, ref. ^13^) and the nuclear deformations caused by DMD mutations^20^, we speculate that this complex may have a structural role in regulating NE morphology and nuclear position in muscle tissue. As multiple members of the dystrophin-glycoprotein complex are transmembrane proteins, it is even possible that the complex penetrates into the perinuclear space, where it may facilitate interactions with proteins of the inner nuclear membrane. In support of this hypothesis, mutations to dystrophin or δ-sarcoglycan alter the distribution of NE proteins lamin A/C, Emerin and and Nesprin-2^20,21^.

### Differential proteomics

Comparing the interactome of a target protein under various conditions can help identify interactions that are important for a specific function. To facilitate such discoveries, we expanded our method to include differential proteomics. By monitoring how a specific treatment or mutation affects the interactome, we identified PPI that are more likely to be relevant to the question at hand. We focused on DNA-PKcs and its regulatory subunits Ku70 and Ku80, and showed that heat shock drives both subunits, but not DNA-PKcs, to associate with lamin A/C.

In HeLa cells expressing progerin or in HGPS fibroblasts, we were able to replicate many known changes to the NE composition, as well as identify novel disease-relevant proteins. These include RanBP2 and Linker of Nucleoskeleton and Cytoskeleton (LINC) complex members nesprin-1 and SUN2, all of which have important roles in laminopathies^22-26^, as well as PTRF, not previously known to associated with progerin. Interestingly, lamin A is known to regulate PTRF transcription^27^ and PTRF and CAV1 inhibit NRF2, promoting stress-induced premature senescence^28^. These proteins show differential abundance in control versus progeria patient-derived fibroblasts, validating our ability to identify clinically relevant targets from patient samples. We note that while PTRF and CAV1 peptides we hardly or not seen in HeLa cells, these proteins gave a robust signal in fibroblasts, as well as primary muscle tissue. Moreover, clear nuclear membrane localization of CAV1 was seen only in muscle samples (Fig. 3J), emphasizing the need to explore changes to the NE composition in a relevant tissue.

### DNA repair and premature aging

Genome wide screens in nematodes and yeast mapped most genes whose downregulation positively affects lifespan, outlining the pathways that regulate aging. However, mutations causing premature aging in mammals have not been well mapped into these pathways. Moreover, the relationships between pathways associated with the known human progeroid syndromes is mostly speculative. One such speculation gives a central role to DNA-PKcs and DNA repair in HGPS premature aging^16^. While no progeroid syndromes resulting from DNA-PKcs mutations have been reported in humans, deletion of DNA-PKcs, Ku70 or Ku80 causes premature aging in mice^29^. Additionally, DNA-PKcs interacts with lamin A/C and the DNA helicase WRN, which when mutated causes Werner’s syndrome. Finally, progeroid features are also observed in conditions resulting from defects in DNA repair, such as Cockayne syndrome and Xeroderma pigmentosum. Our findings support a role for DNA repair pathways in HGPS. We showed that in addition to DNA-PKcs, lamin A/C interacts with Ku70 and Ku80 and this interaction is enhanced when lamin A/C is mutated to progerin leading to potential sequestration of the DNA repair mechanism to the nuclear periphery.

In conclusion, we have developed and implemented a novel method that allows one to identify proteins in proximity to a specific antigen. We used this system to identify proteins in proximity to lamin A/C in various cell lines, primary tissues and patient-derived fibroblasts. Our findings support the hypothesis that the multiple syndromes caused by lamin A/C mutations are driven by tissue-specific interactions.

In future studies, this method can be leveraged to characterize the interactome of various proteins directly from clinical samples of patients and controls, thus accounting for the effect of both genetic and non-genetic factors, like a patient′s life history, on the interactome. Furthermore, BAR can be used to identify interactors of modified proteins or other biomolecules simply by using the appropriate antibody to target the molecule of interest.

## Material and Methods

A list of antibodies, plasmids and cell lines used is presented in Supplementary Table 1^30-32^.

### Biotinylation by Antibody Recognition

A complete protocol is available as a supplementary file. Briefly, samples were fixed in 4% formaldehyde (Thermo Fisher Scientific) for 10-30 minutes at room temperature (RT) and washed with PBST (PBS with 0.1% tween 20). Samples were incubated with 0.5% hydrogen peroxide for 10 minutes to deactivate any endogenous peroxidase activity. Samples were then permeabilized in PBS with 0.5% triton X-100 for 7 minutes and blocked for 2 hours in 1% bovine serum albumin (BSA) in PBST. Samples were incubated with primary antibody overnight, washed with PBST, and incubated with an appropriate secondary antibody conjugated to horseradish peroxidase (HRP) for 3 hours. After extensive washes, samples were incubated with biotin-tyramide (Perkin Elmer) for 10 minutes and a dilution buffer containing hydrogen peroxide was added to a total volume of 150 μL. The reaction was stopped after 1-7 minutes (see Supplementary Dataset) by adding 850 μL of 500 mM sodium ascorbate (Sigma-Aldrich). After two washes with PBST, sample subsets were incubated with FITC-avidin and analyzed under a microscope to validate the expected staining pattern of a given antibody. The remainder of the samples were heated to 99°C for an hour with 1.5% SDS and 1% sodium deoxycholate. Sample volume was adjusted to 1 mL with PBST and biotinylated proteins were extracted with streptavidin beads (Thermo Fisher Scientific) according to the manufacturer′s protocol. For some samples, the presence of a specific protein bound to the beads was validated by Western blot using ∼10% of the beads. Samples were prepared for LC-MS/MS by incubation for 30 minutes at 37°C in 10 mM DTT (Thermo Fisher Scientific) followed by 20 minutes in 50 mM Iodoacetamide (Thermo Fisher Scientific) at room temperature protected from light. Finally, samples were digested overnight in 37°C with 2 μg trypsin (Promega), followed by a second 2 hour digestion.

### Enrichment analysis

For non-SILAC samples, the enrichment factor (E) is set to be min(E_No Ab_, E_Unbound_), where E_No Ab_ is the area ratio of the bound to no antibody control (non-specific binding to beads), and E_Unbound_ is the area ratio of the bound to unbound fractions, normalized to the total area ratios. For proteins found only in the bound sample, E was arbitrarily set to 1000. If the area could be calculated for the bound but not the control sample, E was arbitrarily set to 100. Surprisingly, E was often lower for the target protein than for other known interactors. We speculate this is due to multiple biotinylations, that decreases peptide identification by mass-spectrometry. Known contaminants are shown in the raw data but removed from final lists and analysis. In SILAC samples comparing different mutations or cell lines, protein quantities were normalized, setting the heavy/light lamin A/C ratio to 1.

### Transgene expression

HeLa cells grown in T-75 flasks were transfected using 7 μg per flask of plasmid DNA with lipofectamine 2000 (Thermo Fisher Scientific) in accordance with the manufacturer′s instructions.

### Stable Isotope Labeling by Amino acids in Cell culture

Cells were grown in SILAC MEM medium (Thermo Fisher Scientific) lacking lysine and arginine, supplemented with dialysed 10% FBS (Sigma-Aldrich), 200 mg/l light L-Proline (Sigma-Aldrich) and either heavy L-Lysine (^13^C_6_ ^15^N_2_; 146 mg/l) and L-Arginine (^13^C_6_ ^15^N_4_; 84 mg/l) (Cambridge Isotope Laboratories) or their light equivalents (Sigma-Aldrich). Cells were passaged with non-enzymatic Gibco Cell Dissociation Buffer (Thermo Fisher Scientific). >95% heavy amino acid incorporation rate was validated by mass-spectrometry.

### LC-MS/MS analysis

Protein identification by LC-MS/MS analysis of peptides was performed using an Eksigent nanoLC-Ultra 1D plus system (Dublin, CA) coupled to an LTQ Orbitrap Elite mass spectrometer (Thermo Fisher Scientific, San Jose, CA) using CID fragmentation. Peptides were first loaded onto a Zorbax 300SB-C18 trap column (Agilent, Palo Alto, CA) at a flow rate of 6 μl/min for 6 min, and then separated on a reversed-phase PicoFrit analytical column (New Objective, Woburn, MA) using a 120-min linear gradient of 5–35% acetonitrile in 0.1% formic acid at a flow rate of 250 nl/min. LTQ-Orbitrap Elite settings were as follows: spray voltage 1.5 kV; full MS mass range *m/z* 300 to 2,000. The LTQ-Orbitrap Elite was operated in a data-dependent mode; i.e., one MS1 high resolution (60,000) scan for precursor ions is followed by six data-dependent MS2 scans for precursor ions above a threshold ion count of 500 with collision energy of 35%.

### Database search criteria

Raw files generated by the LTQ Orbitrap Elite were analyzed using Proteome Discoverer v1.4 software (Thermo Fisher Scientific) using Mascot (Matrix Science, London, UK; version 2.5.1) or SEQUEST search engines. The search criteria was set to: database, Swiss Prot (Swiss Institute of Bioinformatics); taxonomy, Human <u>or</u> <u>Mouse</u>; enzyme, trypsin; miscleavages, 2; variable modifications, Oxidation (M), Deamidation (NQ), isotope labeling of lysine (K+8.014 Da) and arginine (R+10.008 Da); fixed modifications, Carbamidomethyl (C); MS peptide tolerance 20 ppm; MS/MS tolerance as 0.8 Da. For the in-gel digestion dataset, identifications were accepted based on one or more unique peptides with a false discovery rate (FDR) of 99% or higher. All other datasets accepted based on two or more unique peptides with a false discovery rate (FDR) of 99% or higher.

### Data analysis

Datasets were imported into a MySQL database. Comparisons involving external datasets was done by converting gene names into the HGNC (http://www.genenames.org/) standard. The mass spectrometry proteomics data have been deposited to the ProteomeXchange Consortium via the PRIDE^33^ partner repository with the dataset identifier PXD004736.

### Human tissue samples

Cadaveric adipose and muscle tissues used in this study were obtained from the National Disease Research Interchange (NDRI) biorepository exempted by the NIH Office of Human Subjects Research.

### Mouse tissue preparation

All animal experiments were approved by NHGRI animal care and use committee. Experiments were performed according to institutional guidelines in accordance with The Guide for the Care and Use of Laboratory Animals and the AVMA guidelines for the Euthanasia of Animals.

C57BL/6 and C57BL/6-TgBAC5-simple-G608G (16492728) mice were housed in a AAALAC Accredited SPF Animal facility.

Perfusion was performed by achieving a surgical plane of anesthesia characterized by loss of consciousness, loss of reflex muscle response and loss of response to noxious stimuli in accordance with the AVMA guidelines for the Euthanasia of Animals using Tribromoethanol (Avertin 1.25%, 0.04 mL/g BW IP). For mass-spectrometry experiments, perfusion of PBS (pH 7.2) followed by 2% Paraformaldehyde in PBS was used for continuous fixation for 60 minutes. Extracted tissues were flash frozen in liquid nitrogen. For electron microscopy, perfusion of PBS followed by 2% Paraformaldehyde or 2% Paraformaldehyde with 0.2% Glutaraldehyde (not compatible with BAR) in PBS was used for continuous fixation for 60 minutes. Tissues were placed in fixation medium for 60 minutes and cryopreserved or directly used.

### Heat shock

T75 Flasks of SILAC labelled HeLa cells were transferred to an incubator preheated to 43°C for two hours and immediately processed.

### Microscopy

Panel 1B was acquired with a ZEISS LSM 880 with Airyscan system equipped with a Plan-Apochromat 63x/1.40 Oil DIC M27 objective and using 488 and 561 nm wavelength lasers. Panels 2A,B S2A Figure S4 were acquired with a DeltaVision PersonalDV (Applied Precision) with either a Plan Apo 60x/1.42 NA or a UPlanSApo 100x/1.4 NA oil lenses. Panels 2G, 3D, S2B and figures S3 and S5 were acquired on a Leica TCS SP5 confocal microscope equipped with a HCX PL APO CS 63.0x/1.40 NA oil lens. FRET experiments were performed using the Leica FRET AB wizard. Panels 3E,G were acquired with a OMX Structured Illumination Super-resolution Scope equipped with a PlanApo 60x/1.40 Oil DIC objective and using 488 and 568 nm wavelength lasers.

## Acknowledgments

We thank Sajni Patel and Marjan Gucek of the NHLBI Proteomics Core for operation of the mass-spectrometer and technical assistance in sample preparation, Stephen M. Wincovitch for microscopy assistance and Julia Fekecs for preparing figure 1A. We thank Lori Bonnycastle and Amanda Dubose for critical reading of the manuscript, useful suggestions and comments.

## Contributions

D.Z.B and F.S.C designed the experiments with input from Y.G., D.Z.B and K.A. performed the experiments, with assistance from M.R.E. U.T. performed the mouse work, D.Z.B analyzed the data. D.Z.B and F.S.C wrote the manuscript, with comments and edits by K.A and Y.G.

## Supplementary information

**Supplementary Figure 1:**
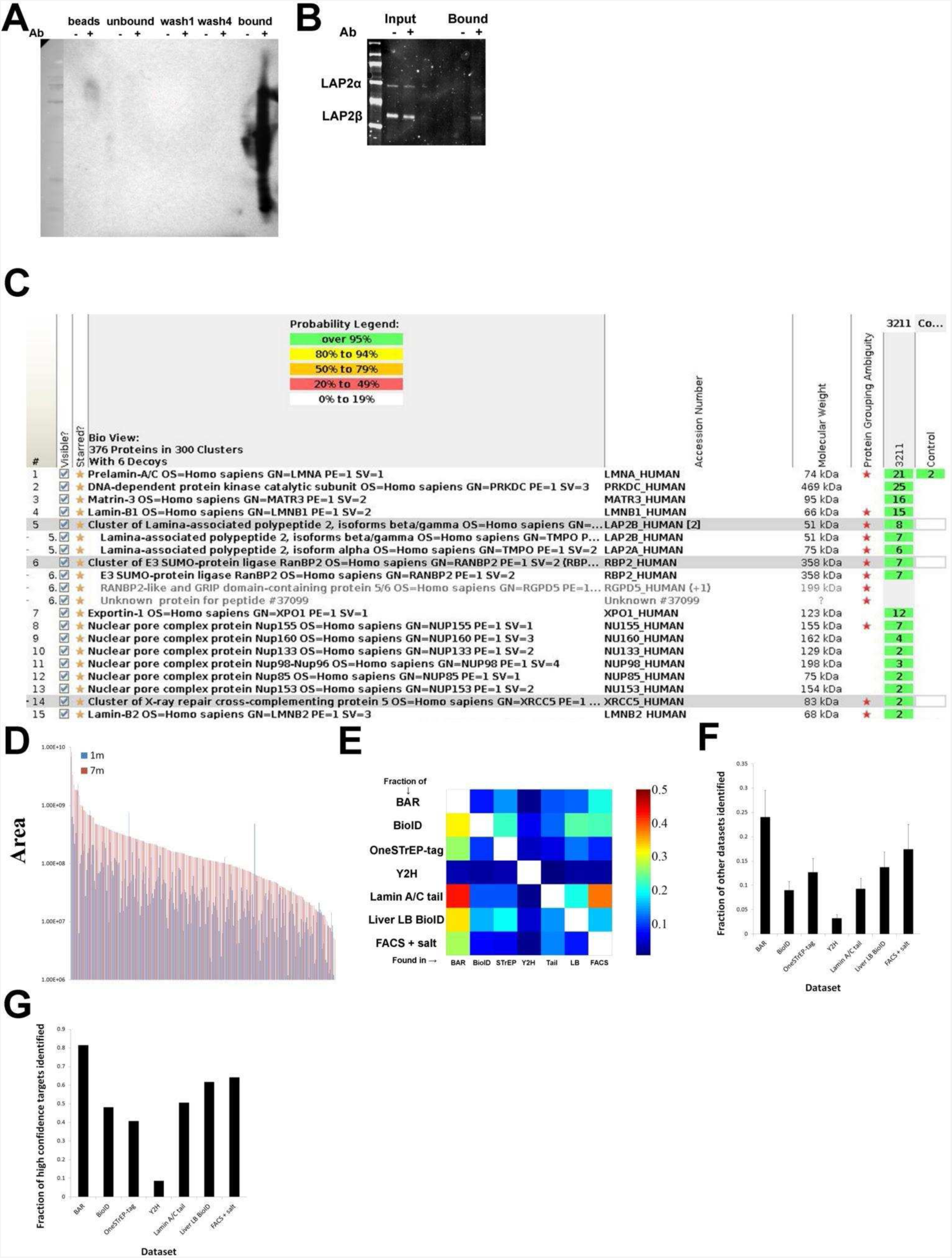
(**A**). Western blot using Strepavidin-HRP showing antibody guided efficient biotin labeling and recovery of proteins. (**B**) LAP2, a known lamin A/C binding protein, is pulled down by BAR, as evident from a Western blot. (**C**) A scaffold (Proteome Software, Portland, Oregon) screenshot showing multiple known NE and lamin A/C interacting proteins, and their peptide counts in the antibody sample and control. (**D**) Signal intensity of 1 minute in-gel digestion vs 7 minutes on bead digestion. (**E**) Heat-map showing the fraction of each dataset covered by any other dataset. (**F**) For each of the 7 datasets (our in-gel digestion dataset, as well as 6 other published datasets), the average fraction of all other datasets covered by it is shown. (**G**) The fraction of high confidence targets identified in each dataset is shown.

**Supplementary Figure 2:**
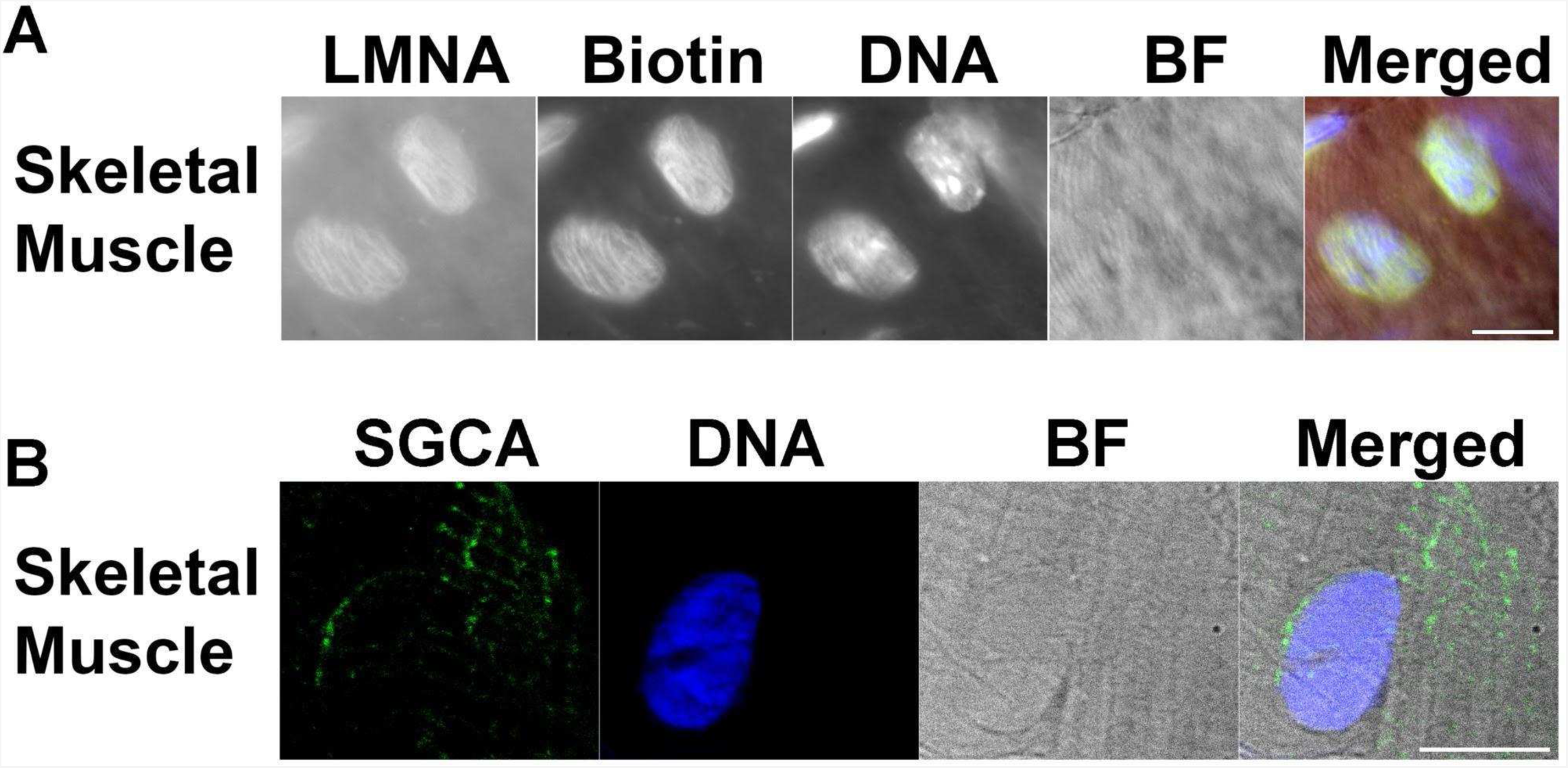
(**A**). Primary Mouse skeletal muscle stained for lamin A/C. BF - brightfield. (**B**) Primary human skeletal muscle tissue showing SGCA staining at the nuclear periphery.

**Supplementary Figure 3:**
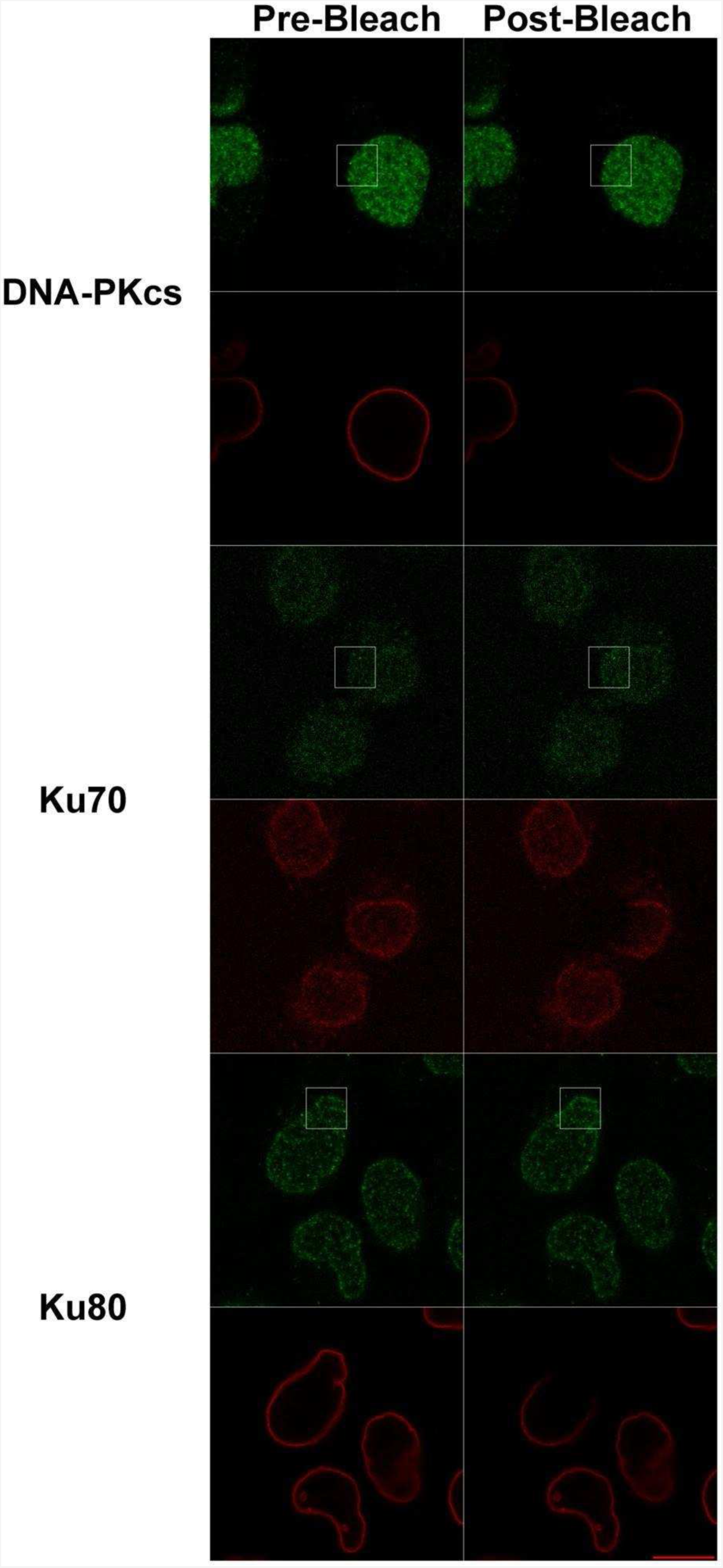
Förster resonance energy transfer (FRET) show that for Ku70 and Ku80, but not DNA-PKcs, fluorescence intensity increases after bleaching of lamin A/C adjacent fluorophores. FRET efficiency was 0 for DNA-PKcs, 0.34 for Ku70 and 0.15 for Ku80. We note that negative FRET results, as is the case for DNA-PKcs, cannot be interpreted as a lack of interaction.

**Supplementary Figure 4:**
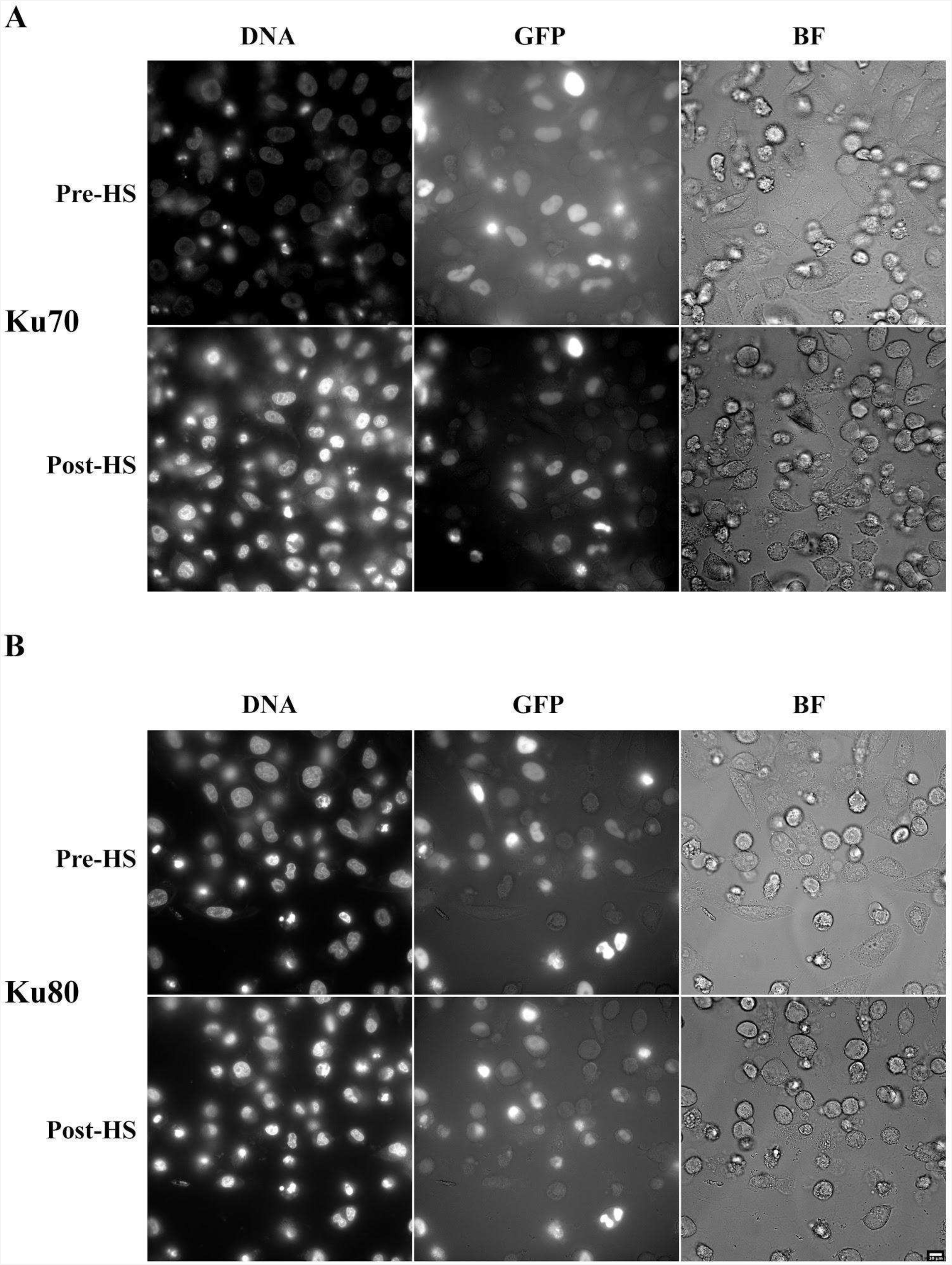
Pre and post heat shock images of Hela cells expressing (**A**) Ku70-GFP or (**B**) Ku80-GFP.

**Supplementary Figure 5:**
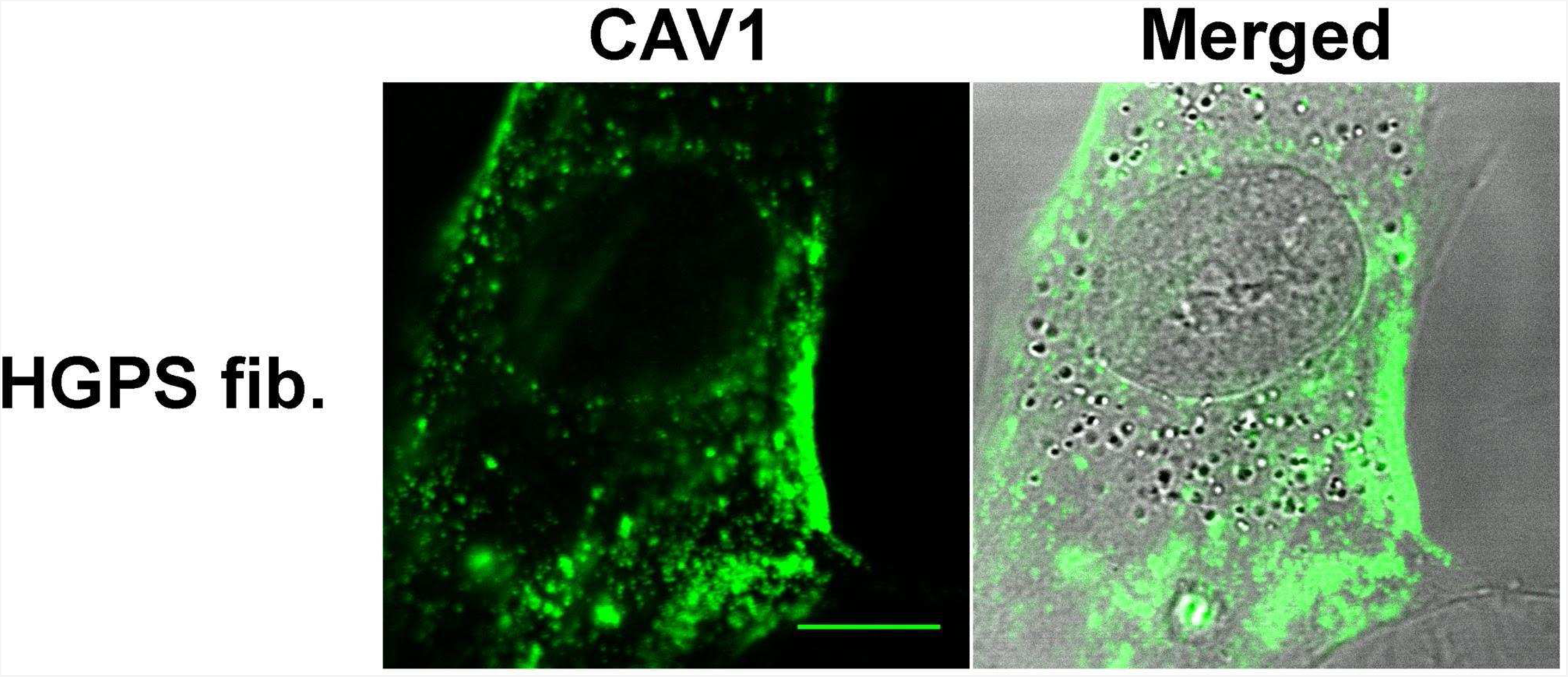
HGPS fibroblasts stained for CAV1 and merged with brightfield.

